# Stress granule induction in rat retinas damaged by constant LED light

**DOI:** 10.1101/2024.04.26.591385

**Authors:** María M. Benedetto, Melisa Malcolm, Manuel G. Bruera, Laura G. Penazzi, Mario E. Guido, María A. Contín, Eduardo Garbarino-Pico

## Abstract

**Objectives:** Stress granules (SGs) are cytoplasmic biocondensates formed in response to various cellular stressors, contributing to cell survival. While implicated in diverse pathologies, their role in retinal degeneration (RD) remain unclear. We aimed to investigate SG formation in the retina and its induction by excessive LED light in a RD model.

**Methods:** Rat retinas were immunohistochemically analyzed for SG markers G3BP1 and eIF3, and SGs were also visualized by RNA FISH. Additionally, SGs were induced in primary retinal cell and eyeball cultures using sodium arsenite. Light exposure experiments utilized LED lamps with a color temperature of 5,500 K and 200 lux intensity for short-term or 2-8-day exposures.

**Results:** SGs were predominantly detected in retinal ganglion cells (RGCs) and inner nuclear layer (INL) cells, confirmed by sodium arsenite induction. SG abundance was higher in animals exposed to light for 2-8 days compared to light/dark cycle controls. RGCs consistently exhibited more SGs than INL cells, and INL cells more than outer nuclear layer cells (Scheirer-Ray-Hare test: H 13.2, p = 0.0103 for light condition, and H 278.2, p < 0.00001 for retinal layer). These observations were consistent across four independent experiments, each with three animals per light condition.

**Conclusions:** This study identifies SGs in the mammalian retina for the first time, with increased prevalence following excessive LED light exposure. RGCs and INL cells showed heightened SG formation, suggesting a potential protective mechanism against photodamage. Further investigations are warranted to elucidate SGs’ role in shielding against light stress and their implications in retinopathies.

## INTRODUCTION

Retinal degeneration (RD) is a neurodegenerative disease with numerous contributing factors, encompassing processes like remodeling, photoreceptor death, and the deterioration of both the structure and function of this tissue^1^. The eye has evolved protective mechanisms to safeguard against excessive light exposure, which can be detrimental to the retina^2,3^. Artificial light alters the natural illumination, leading to adverse effects on retinal function. Light-emitting diodes (LEDs) have become the primary source of both household and public lighting. They are also integral to modern technologies like computers, TVs, tablets, cell phones, and gaming consoles. As a result, our visual system is expose to LED light in excess. Despite the cost-effectiveness and energy efficiency of LEDs, they emit a significant amount of blue light, with wavelengths ranging from 460 to 500 nm^4^. This blue light can potentially have detrimental effects on human vision^5^.

The retina is the most energetically demanding tissue, known for its abundant oxygen supply and rich content of polyunsaturated fatty acids. It also contains elevated levels of photosensitizers, further rendering it susceptibility to oxidative stress^6–8^. The production of reactive oxygen species (ROS) in the context of oxidative stress is a crucial component of the common pathway leading to neural damage in various acute and chronic neurological eye disorders^9^. While ROS play important roles in cell signaling and regulation, their excessive production can result in damage to cellular macromolecules, including DNA, proteins, and lipids^10^.

We have previously established a model of RD induced by continuous exposure to low-intensity LED light (LL) in Wistar albino rats^11^. Our research has revealed an elevation of ROS production within the outer nuclear layer (ONL) of retinas from animals exposed to LL for 5 days. Concurrently, there has been a decline in docosahexaenoic acid (DHA), a crucial component of rod cell external segment membranes, likely due to oxidative processes^12^. Additionally, significant photoreceptor cell loss has been observe after 7 days of LL exposure, accompanied by heightened rhodopsin phosphorylation early in the exposure period. Notably, most retinal ganglion cells (RGCs) and inner nuclear layer (INL) cells remain viable, albeit with alterations in the expression and localization of photopigments like melanopsin (OPN4) and neuropsin (OPN5)^13^. These findings underscore the varied susceptibility of distinct retinal cell types to damage induced by excessive LED light exposure. This heterogeneity is anticipated given the diverse array and distribution of photopigments across retinal cells, along with alternative phototransduction pathways. Moreover, the observed differences may also stem from the presence of distinct defense mechanisms against photodamage among these cell types.

Cellular stress triggers an adaptive program known as the integrated stress response (ISR) enables to endure and survive. Nevertheless, if cells are unable to overcome it, it can also activate the cell death mechanism to eliminate damaged cells^14^. Depending on the intensity and duration of the stress, one of these two responses prevails. A prominent feature of the ISR is its ability to inhibit global protein synthesis promoting the accumulation of translationally stalled 48S complexes that undergo liquid-liquid phase separation. These complexes form biocondensates of RNA and proteins known as stress granules (SGs)^15,16^. SGs are enriched in polyadenylated mRNA, small ribosomal subunits, translation initiation factors, RNA-binding proteins (RBPs), and other factors^17–19^. The function of SGs has been linked to the regulation of translation, stability and storage of cytoplasmic mRNA, however this is not completely elucidated and is a matter of controversy^20^. Considering that many molecules linked to signaling pathways are concentrated in the SG, it has been proposed that they act as signaling hubs^21^. Numerous factors linked to apoptosis are concentrated in SG, inhibiting several pro-apoptotic signaling pathways and favoring cell survival^22–27^. The formation of SGs also diminishes the accumulation of ROS^28^. The aim of this study was to characterize the formation of SGs in the retina and assess their prevalence in retinal light damage.

## MATERIALS AND METHODS

### Animals

All procedures conducted adhered to the guidelines outlined in the ARVO statement for the use of animals. Additionally, all protocols were approved by the local animal committee (School of Chemistry, UNC, Exp. EXP#2023-00453889-UNC-ME#FCQ). Male albino Wistar rats, aged 12-15 weeks, bred in our laboratory for five years, were housed under a 12-hour light/12-hour dark (LD) cycle. White fluorescent light of ∼50 lux intensity was provided. Throughout the experiment, the rats had *ad libitum* access to food and water.

### Immunohistochemistry (IHC)

IHC was conducted as before^11^. Rat eyes were fixed in 4% (w/v) paraformaldehyde in PBS overnight (ON) at 4°C, cryoprotected in sucrose 30% (w/v) and mounted in an optimal cutting temperature compound (OCT; Tissue-Tek^®^, Sakura). Then, 20 µm retinal sections were cut along the horizontal meridian (nasal-temporal) by cryostat (HM525 NX-Thermo Scientific). Sections were washed in PBS and permeabilized with PBS-T (PBS with 0.5% Triton X-100), for 90 min at room temperature (RT). Then, they were blocked with blocking buffer [PBS supplemented with 3% (w/v) BSA, 0.1% (v/v) Tween 20, 1% (v/v) Glycine and 0.02% (w/v) Sodium Azide] for 2.5 h at RT and incubated with anti-eIF3 (1:300, Santa Cruz sc-16377) or anti-G3BP1 (1:1000, BETHYL Laboratories A302-033A), two robust SGs markers^15,29–31^, diluted in blocking buffer, ON at 4°C in a humidified chamber. Samples were then rinsed in PBS-TW (0.1% (v/v) Tween 20 in PBS) and incubated with goat anti-rabbit IgG Alexa Fluor 488 or 549 (1:1000) respectively and DAPI (3 μM), for 1 h at RT. Finally, they were washed 3 times in PBS and mounted with Mowiol (Sigma-Aldrich).

### Fluorescence *in situ* hybridization (FISH) and immunofluorescence (IF)

Samples were obtained and fixed as detailed in the preceding section. FISH-IF was carried out according to Meyer and colleagues^32^. Fixed samples were permeabilized with TBS-T ( 0.01 M Tris buffer pH 7.4, 0.1M NaCl, 0.2% (v/v) Triton X-100), 20 min at RT, washed 5 min with TBS and pre-hybridized using hybridization buffer (H-7140, Sigma-Aldrich) for 1 min at RT. Subsequently, they were hybridized with Cy3-Oligo(dT)_30_ (Sigma-Aldrich) diluted in hybridization buffer, at a final concentration of 20 nM, ON at 40 °C in a humidified chamber. Then, samples were rinsed once with Washing Buffer 1 (50% (v/v) Formamide 12.6 M, 0.25M NaCl, 0.075 M Tris Buffer (pH 8.5), and 0.1% (v/v) Tween 20) 5 min at RT in constant shaking, four times with Washing Buffer 2 (0.05M NaCl, 0.075 M Tris Buffer (pH 8.5) and 0.1% (v/v) Tween 20) for 15 min each in constant shaking and with TBS 5 min. Then, they were blocked with blocking buffer for 2.5 h at RT with continuous shaking and incubated ON with anti-eIF3 (Santa Cruz sc-16377) diluted in blocking buffer at 4°C in a humidified chamber. Samples were then rinsed in TBS and incubated with goat anti-rabbit IgG Alexa Fluor 488 (1:1000) and DAPI (3 μM), for 1 h at RT. Finally, they were washed 3 times with TBS-T for 5 min each, once with TBS for 5 min and mounted in Mowiol.

### Primary Cell cultures

Primary cultures were obtained from rats at post-natal day 7^33^. Retinas were manually dissected with gentle up and down passes in ice-cold Ca^+2^–Mg^+2^-free Tyrode’s buffer and treated with papain (P3125, Sigma–Aldrich) 20 min and deoxyribonuclease I (18047-019, Invitrogen) 10 min at 37°C under constant air flow in a humid atmosphere. Cells were precipitated at 5000g for 20 min at 4°C and resuspended in 10% fetal bovine serum (FBS, Gibco)-DMEM. Cells were seeded in coverslips treated with poly-L-lysine (10 μg/mL) and grown for 6 days in Neurobasal medium (Gibco) supplemented with 0.05% (v/v) Amphotericin B, 0.1% (v/v) Forskolin (Sigma Aldrich), 0.02% (v/v) Recombinant Human BDNF (R&D Systems), 2% (v/v) B27 (GIBCO), and 1% (v/v) L-glutamine (Glutamax, Gibco).

### SG induction by sodium arsenite

To induce SG formation in cell cultures, sodium arsenite (NaAsO_2_, Sigma-Aldrich S7400) was added to the media at final concentrations of 250 or 500 μM and incubated for 30 minutes. For *ex vivo* retinal SG induction, eyeballs were dissected and subjected to intravitreal injection with 2 μl of 50 mM arsenite or vehicle (sterile MQ H_2_O), followed by a 30-minute incubation at 37°C in 10% FBS-DMEM. Subsequently, the eyes were fixed and processed for IHC.

### Immunocytochemistry (ICC)

Cells were washed with PBS, fixed 15 min in 4% PFA and permeabilized 10 min with −20°C methanol at RT, according to Kedersha & Anderson^31^. After 3X-washes with cold PBS for 5 min, they were incubated for 1 h with blocking buffer. Then, samples were incubated with anti-G3BP1 (1:1000, A302-033A, BETHYL Laboratories) and anti-DM1A (1:1000, Sigma) diluted in blocking buffer, ON at 4°C in a humidified chamber. Samples were then rinsed in PBS-T and incubated with goat anti-rabbit IgG Alexa Fluor 488 or 549 (1:1000) respectively and DAPI (3 μM), for 1 h at RT. Finally, they were washed 3 times with PBS and mounted with Mowiol.

### Light exposure protocols and induction of retinal degeneration (RD)

RD was induced according Contín et al.^11^. Animals were exposed to constant light (LL) for varying durations: 54 h (LL2, ∼2 days), 102 h (LL4, ∼4 days), 150 h (LL6, ∼6 days), and 198 h (LL8, ∼8 days). LED lamps (EVERLIGHT Electronic Co., Ltd. T-13/4 3294-15/T2C9-1HMB, color temperature of 5,500 K) were installed in specially designed boxes, with the temperature-controlled at 24 ± 1°C. Light intensity at the level of the rats’ eyes was measured at 200 lux using a light meter (model 401036; Extech Instruments Corp., Waltham). For short-duration light exposure experiments, identical stimulation boxes were utilized for intervals of 0-12 h. Control retinas were obtained from animals maintained in an LD cycle (50 lux white fluorescent light), dissected 6 h after the lights were turned on. Euthanasia was performed using a CO_2_ chamber.

### Image acquisition and quantification of SGs

Confocal imaging was conducted using a FluoView FV1200 confocal microscope (Olympus). A 60x/1.3 silicone immersion objective (UPLSAPO60XS, Olympus) captured images at a resolution of 2048×720 pixels. Image Z stacks were obtained with a pinhole of adjusted for 2 µm optical slices and 1.4 zoom, Kalman 0. ImageJ software was used for image processing.

For SG quantification, each slice within the Z stacks underwent the following procedure: 1) Background subtraction (value=70), Outlier Removal (radius=1, threshold=200), Background Rolling Ball Radius Subtraction (radius=3), and Gamma adjustment (value=2). The stack was duplicated, and Gaussian filters with radii of 1 (g1) and 3 (g3) were applied, followed by subtraction (g1-g3). A maximum intensity projection was created from the resulting stack. 2) Manual selection of each retinal layer area. 3) Counting SGs within each ROI corresponding to the layers using Process > Find Maxima > Prominence = 100-200. Quantification of GCL and INL nuclei was manual. ONL nuclei quantification employed Process > Find Maxima > using a prominence value to select each nucleus only once.

### Statistical Analysis

The analysis was conducted with RStudio. Normality and homogeneity of the variance assumptions were evaluated using the Shapiro-Wilks and the Bartlett tests, respectively. The non-parametric Scheirer-Ray-Hare test was used to verify the significance difference between the days of light treatment and between layers of the retina. In case there was a significant difference, the Dunn‘s ad hoc test with a Bonferroni correction were employed. We also compared the number of SGs formed in the different retinal layers throughout the light treatment. The Kruskal-Wallis test was used to verify if there was a statistically significant difference between treatments in INL and ONL, followed by Dunn‘s Test with a Bonferroni correction. GCL was analyzed using an ANOVA test followed by Tukey’s post hoc test. Similarly, for each light treatment condition, a non-parametric Kruskal-Wallis test followed by an ad hoc Dunn‘s test with a Bonferroni correction were performed to see if there was a difference between the layers of the retina.

### Manuscript writing

The authors initially drafted the manuscript, after which ChatGPT 3.5 was asked to enhance the English, and then the authors reviewed it once again.

## RESULTS

### Detection and characterization of SGs in rat retina

To characterize SGs within the retina, we initially performed IHC using antibodies targeting the established SG marker G3BP1^15,30,31^. Figures 1A and 1B illustrates the G3BP1 immunolabeling in retinas exposed for 48 h to 200 lux LED light. Numerous distinct spots resembling SGs are evident, consistent with their typical appearance. Given the protein’s localization within both these biocondensates and the cytoplasm, a background signal is also observable. Predominantly, these foci are observed within the RGCs, with some presence in the INL and rare occurrence in the ONL.

**Figure 1:**
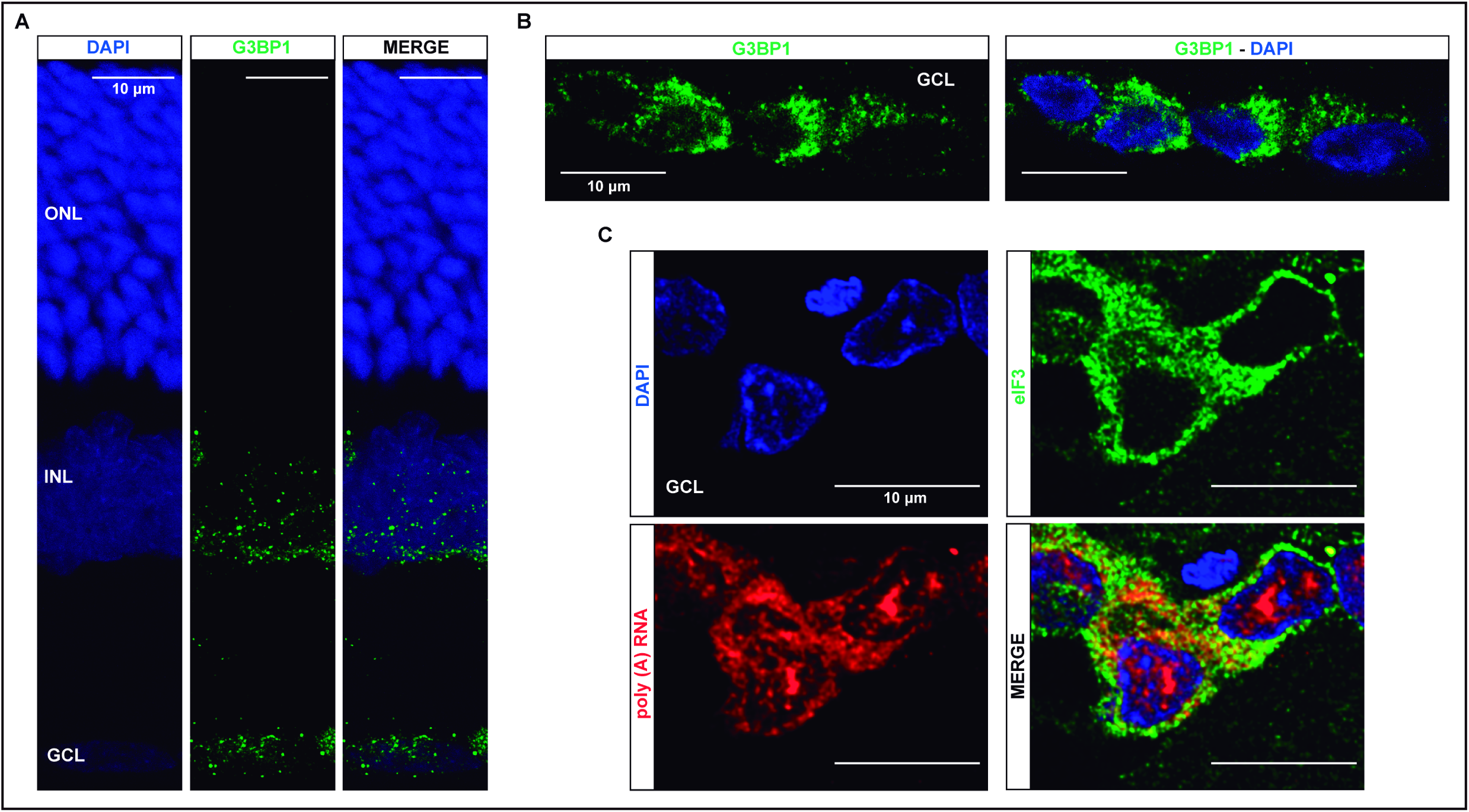
Visualization of Stress Granules (SGs) in Rat Retina. **A.** Immunohistochemical (IHC) staining of retinas from rats exposed to constant LED light (200 lux intensity) for 2 days (LL2). Images acquired by confocal fluorescence microscopy depict the SG marker G3BP1 detected using specific antibodies. Nuclei stained with DAPI (blue); G3BP1 visualized in red. Scale Bar: 10 μm. **B.** Higher-resolution images of retinal ganglion cells (RGCs) from experiments conducted similarly to panel A. **C.** Simultaneous analysis of RGCs using poly(A)+ RNA fluorescence in situ hybridization (Poly(A)+ RNA-FISH) and immunofluorescence (IF). Cy3-Oligo(dT)30 probe binds to the 3’-poly(A) end of polyadenylated mRNAs concentrated within SGs. Anti-eIF3 antibody, another SG marker, was used. Nuclei stained with DAPI (blue); eIF3 in green; Poly(A)+ RNA in red. Scale Bar: 5μm. (See Materials and Methods for experimental details).

To further confirm the presence of SGs in the retina, we conducted dual-labeling of SGs using Poly(A)+ RNA Fluorescent In Situ Hybridization (Poly(A)+ RNA-FISH) in combination with immunofluorescence utilizing an antibody targeting eIF3, another established SG marker^15,29,31^. Cy3-Oligo(dT)_30_, a probe specifically binding to the poly(A) tail of mRNA concentrated within SGs, was employed. Figure 1C shows the co-localization of both signals, Poly(A)+ RNA and eIF3, within bright spots, providing evidence for the presence of SGs within retinal tissue, primarily concentrated in RGCs.

### Induction of retinal SGs by sodium arsenite

To validate the identity of the structures identified, we conducted experiments to determine whether sodium arsenite, a well-established inducer of SGs^31,34,35^, could effectively augment the quantity and/or size of the observed bright spots detected by IHC. In Figure 2A, ICC demonstrates G3BP1 labeling in primary cell cultures of the total rat retina. Non-treated cells (control) display a diffuse cytoplasmic labeling of G3BP1. In contrast, cells treated with NaAsO_2_ exhibit distinct dot-shaped structures characteristic of G3BP1 concentration within SGs. Notably, a higher number of these distinct dot-shaped structures were visible at the higher arsenite concentration utilized.

**Figure 2:**
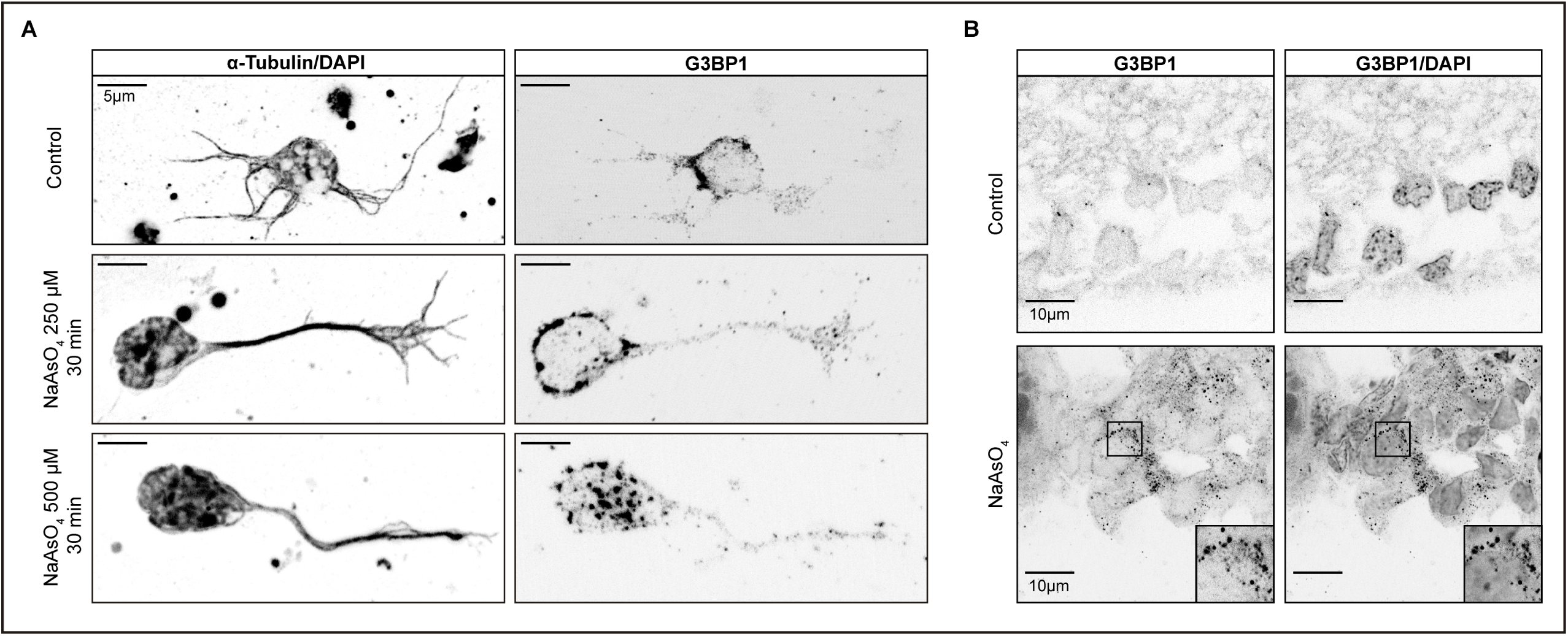
Induction of Stress Granules with arsenite in cultured cells and retinal tissue. **A.** Primary retinal cultures from rats at post-natal day 7 were treated with the SG inducer sodium arsenite at concentrations of 250 or 500 μM for 30 minutes, or left untreated as control. The left panels display α-Tubulin and DAPI signals, while the right panels show G3BP1 staining. Scale Bar: 5 μm. **B.** Ocular globes were dissected and immediately intravitrealy injected with 2 μl of 50 mM sodium arsenite or vehicle solution (sterile MQ H_2_O), followed by a 30-minute incubation at 37°C in 10% FBS-DMEM. Subsequently, the eyes were fixed and processed for IHC. The left panels show the signal from G3BP1, and the right panels show G3BP1 and DAPI staining. Scale Bar: 10 μm.

Subsequently, we investigated the impact of arsenite on *ex vivo* retinal tissue. Eyecups were dissected, subjected to intravitreal injection with arsenite, and cultured for 30 minutes prior to IHC analysis. In Figure 2B, a few discernible bright spots are evident in non-treated RGCs (control); however, a marked rise in SG-like dots is observed in RGCs from retinas treated with NaAsO_2_. Remarkably, the highest count of bright spots was observed in RGCs. These findings substantiate that treatment with NaAsO_2_ induces the formation of SGs, both in cell culture and *ex-vivo* retinal experiments.

### Stress Granules (SGs) in Retinas Exposed to Short-Term Low-Intensity LED Light

In Figure 1, we detected SGs in animals exposed to constant light for 48 h, here, we aimed to ascertain if shorter durations (0-12 h) of low-intensity LED light exposure (200 lux) could induce the formation of these cellular biocondensates. Figure 3 illustrates the presence of these granules in both control conditions and at three distinct time points following light exposure. However, the differences observed among the various time intervals were not statistically significant.

**Figure 3:**
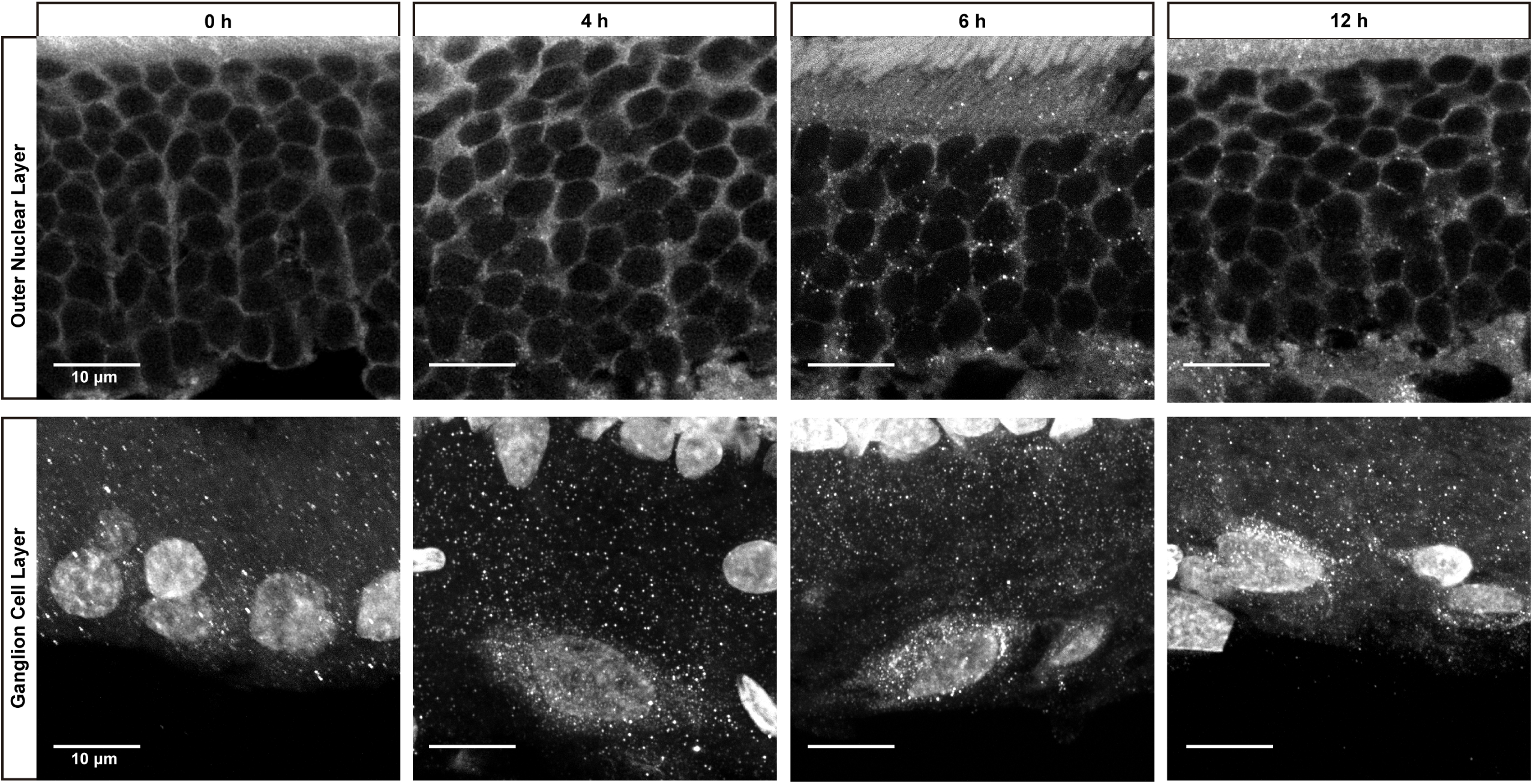
Stress Granules in retinas exposed to short periods of low-intensity LED light. Rats, adapted to a 12 h light-12 h dark (LD) cycle (white fluorescent light <50 lux), were subjected to light exposure experiments. 6 h after the onset of the light phase of the LD cycle, rats were exposed to 200 lux intensity LED light for 0 (control), 4, 6, or 12 h. Following exposure, retinas were dissected and subjected to IHC using an anti-G3BP1 antibody to visualize SGs. Representative images were captured from outer nuclear layer (ONL) cells (upper panel) and RGCs (lower panel). Nuclei were stained with DAPI, while SGs were labeled with anti-G3BP1 antibody. Scale bar: 10 µm.

### SG Formation in Rat Retinas Exposed to Prolonged Low-Intensity LED Light

To examine SGs within our RD model —a condition where animals are continually exposed to 200 lux LED light for 2-8 days— we assessed SG counts across various retinal layers in control conditions and during the progression of RD. Our objective was to pinpoint the particular retinal cell types exhibiting heightened SG formation in response to prolonged light exposure. Figure 4A shows representative retinal images in which SGs were labeled with anti-G3BP1. In order to quantify SGs as a function of light exposure time, we analyzed retinas of animals exposed to 2-8 days of LL. Four independent experiments were carried out, in each of them 3 animals per group were used and 5-7 images were taken for each lighting condition (n=20-28/group). Figs. 4B-C and Table 1 show the results of the four experiments analyzed together. To analyze the combined effect of the studied factors, we employed the Scheirer-Ray-Hare test. Both light exposure time (H=13.2; p=0.01034) and retinal layer (H=278.2; p<0.00001) exhibited significant changes in SG numbers when normalized by the number of cells (total nuclei of each layer). No combined effect of both factors was observed (H=3.301; p=0.91408). *Post-hoc* analysis (Dunn’s test with Bonferroni correction) revealed lower SG counts per cell in control retinas compared to those of animals maintained for 2, 6, or 8 days in LL (Fig. 4B). In all cases except LD controls, SG numbers were significantly higher in RGCs than in INL cells, and higher in INL cells than in ONL cells, where SGs were rare (Fig. 4, Table 1 and Supplementary Material 1). Similar results were obtained when each factor was analyzed separately using the Kruskal-Wallis rank sum test (Fig. 4C-E, Supplementary Material 1). Considering the variability in cell number and size within each layer, we further normalized the data by area (Supplementary Material 2). The results remained consistent, with a greater number of SGs in light-treated retinas, and RGCs harboring more SGs than INL cells, which, in turn, contained more SGs than ONL cells.

**Figure 4:**
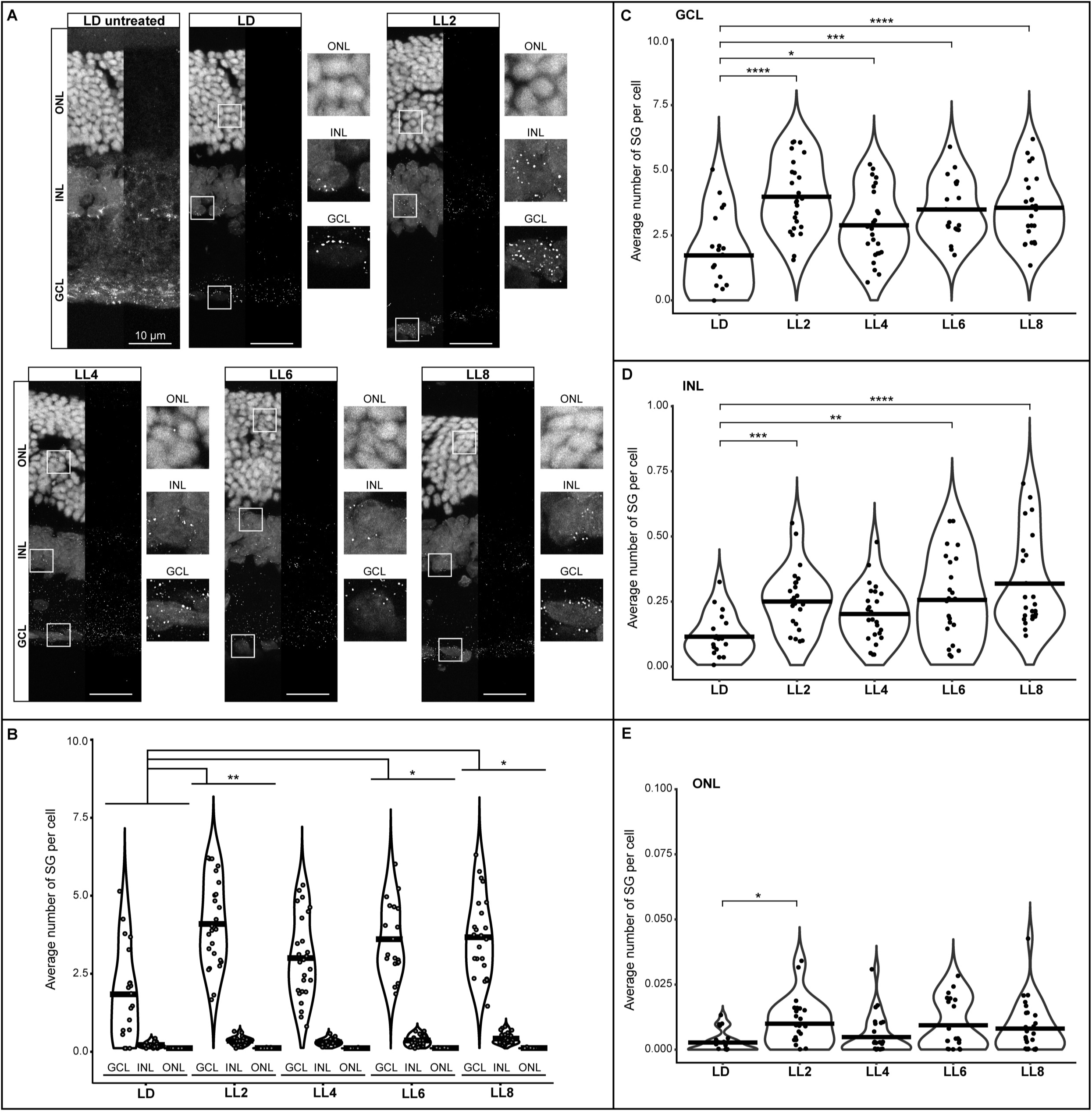
Induction of Stress Granules in the inner retina by excessive low-intensity LED light exposure in a model of Retinal Degeneration. Animals were exposed to constant LED light (200 lux, color temperature of 5,500 K) for 54 h (LL2, ∼2 days), 102 h (LL4, ∼4 days), 150 h (LL6, ∼6 days), and 198 h (LL8, ∼8 days), in temperature-controlled chambers at 24 ± 1°C. Control retinas were obtained from animals maintained in a LD cycle (white fluorescent light <50 lux) and dissected 6 h after the lights were turned on. **A.** Representative images of retinas analyzed by IHC using an anti-G3BP1 antibody. LD untreated and LD represent the same image before and after processing for quantification (See M&M). All other images are shown after processing. Scale bar: 10 µm. **B.** Quantification of SGs per cell in different layers of the retina in each light condition. Data represent results from four independent experiments, each using three animals per group, with 4–6 images acquired per case (n = 16-24 images per group). The Scheirer-Ray-Hare test was used for analyzing all light conditions and retinal layers together (H = 13.2, p = 0.01034 for light conditions; H = 278.2, p < 0.00001 for retinal layer; H = 3.301, p = 0.91408 for interactions between the two factors; see Table 1 and Supplementary Table 1). The results from LL2, LL6, and LL8 were higher than LD (Dunn’s test with a Bonferroni correction). **C-E.** Data from the GCL (C), INL (D), and ONL (E) are visualized for comparison within each retinal layer. GCL was analyzed by ANOVA test followed by Tukey’s post hoc, whereas INL and ONL were analyzed by Kruskal-Wallis rank sum test followed by Dunn’s test with a Bonferroni correction. Significance is indicated by asterisks ( * p < 0.05, ** p < 0.01, *** p < 0.001, **** p < 0.0001).

**Table 1:** Number of SGs per cell.

|  | GCL |  | INL |  | ONL |  |
| --- | --- | --- | --- | --- | --- | --- |
|  | Mean | SD | Mean | SD | Mean | SD |
| LD | 1.73 | 1.51 | 0.11 | 0.09 | 0.00 | 0.00 |
| LL2 | 3.99 | 1.38 | 0.24 | 0.13 | 0.01 | 0.01 |
| LL4 | 2.89 | 1.33 | 0.20 | 0.11 | 0.00 | 0.01 |
| LL6 | 3.49 | 1.17 | 0.25 | 0.17 | 0.01 | 0.01 |
| LL8 | 3.56 | 1.29 | 0.31 | 0.18 | 0.01 | 0.01 |
| Mean in LL | 3.48 |  | 0.26 |  | 0.01 |  |

## DISCUSSION

In this study, we employed a multi-faceted approach, utilizing IHC with antibodies targeting two SG markers, eIF3 and G3BP1, along with dual-labeling through Poly(A)+ RNA-FISH and immunofluorescence with an eIF3-specific antibody. Through these methodologies, we successfully identified SGs in the rat retina. To validate the authenticity of our SG detection, we conducted experiments using sodium arsenite, a well-established inducer of SG formation. Remarkably, retinas or retinal cell cultures treated with arsenite exhibited a significant increase in SG-like dots compared to control retinas, which showed minimal SG presence. Furthermore, we investigated SG development in a model of RD induced by prolonged exposure to constant low-intensity LED light. After 48 hours of light exposure, a notable increase in SGs was observed in both RGCs and the INL, indicating that excessive light exposure indeed induces SG formation. Notably, the layers of the retina that demonstrated higher SG quantities were the ones that exhibited greater resilience in the face of RD, highlighting a potential link between SG formation and cellular survival in this degenerative model.

The majority of investigations into SGs have been conducted in cell cultures, often induced under non-physiological conditions. While SG presence has been documented in neurons, such as in mouse primary cortical neurons^36^, we observe a dearth of prior descriptions regarding SGs in the retina. Our search yielded only one account of G3BP1-containing cytoplasmic granules in the rat retinal ganglion cell line RGC-5^37^. Notably, uncertainties persist concerning the origin and nature of this cell line^38–41^. Although SGs have been identified in cultured retinal pigment epithelium cells^42^, our study constitutes the inaugural documentation of SGs in the retina and their response to a environmental stimulus, such as light.

Considering the pro-survival role attributed to SGs and our finding that they are induced in the inner retina in response to prolonged exposure to low-intensity LED light, it can be hypothesized that these granules play a role in the survival of RGCs and INL cells. In these cells, the number of SGs increases, whereas in cones and rods, the neurons that undergo cell death, are scarcely distinguishable. In this study, we observed a correlation, and future experiments are required to determine the validity of this hypothesis.

Conversely, SGs have also been associated with neurodegenerative diseases such as amyotrophic lateral sclerosis (ALS), frontotemporal dementias (FTD), and Alzheimer’s disease (AD)^16^. Some proteins associated with SGs have been linked to the formation of pathological protein aggregates (e.g., TDP-43 and FUS proteins). In many cases, the formation of these aggregates is associated with mutations in proteins normally found in SGs, which, when modified, impact the disassembly and clearance of the granules^16,43^.

In our approach, we cannot distinguish whether the detected SGs favor cell survival or, conversely, in the context of continuous light exposure, chronically promote the possibility of producing pathological aggregates. In the majority of experimental paradigms used to study SGs, their induction occurs under conditions of acute stress, typically confined to a short duration of minutes or a few hours. In the model we employed, the stress is chronic (2-8 days). We are unaware of the dynamics of SG formation and clearance. For instance, SGs could initially form rapidly in photoreceptor cells, which are most susceptible to light stress, but unable to survive, triggering death processes that do not involve SG formation.

Our study reveals, for the first time, the presence of SGs in the mammalian retina, with their numbers increasing in response to excessive LED light exposure. Significantly, their prevalence is notably higher in inner retinal cells, which exhibit a remarkable resistance to light-induced damage. Further investigations are necessary to determine whether SGs play a pivotal role in shielding against light stress or potentially contribute to other retinopathies.

## ACKNOWLEDGMENTS

The authors express their gratitude to Dr. Cecilia Sampedro, Dr. Carlos R. Mas, and Dr. Alejandra Trenchi for their invaluable technical support in image acquisition and analysis. Special thanks are extended to Rosa Andrada for her exceptional management of the animal facility. This work was made possible by the generous support of grants from the Agencia Nacional de Promoción Científica y Técnica (PICT 2020 No. 02699), Consejo Nacional de Investigaciones Científicas y Tecnológicas de la República Argentina (CONICET PIP 2020), Secretaría de Ciencia y Tecnología de la Universidad Nacional de Córdoba (SeCyT-UNC), and the Ministry of Sciences and Technology of Córdoba.

