## Supplemental material 1 for "Stress granule induction in rat retinas damaged by constant LED light"

### Supplemental Material 1: Statistical analysis of SG number per cell changes in retinal layers during DR progression (Fig. 4B-C)

#### a) Combined effect of light exposure duration and retinal layer on SG count per cell

**Scheirer-Ray-Hare test**

| | |
|---|---|
| **DV** | Number SGs / cell |
| **Observations** | 359 |
| **D** | 1.9961031 |
| **MS total** | 10711.17 |

| | Df | Sum Sq | H | p |
|---|---|---|---|---|
| **Light treatment** | 4 | 140824 | 13.2 | 0.01034 |
| **Retinal layer** | 2 | 2968178 | 278.2 | 0.00000 |
| **Light:Layer** | 8 | 35216 | 3.301 | 0.91408 |
| **Residuals** | 342 | 655509 | | |

**Dunn`s test with a Bonferroni correction: light treatment (Z and p)**

| | LD | LL2 | LL4 | LL6 |
|---|---|---|---|---|
| **LL2** | -3.3693<br>**0.0038** | | | |
| **LL4** | -2.3330<br>0.0982 | 1.1227<br>1.0000 | | |
| **LL6** | -2.8090<br>**0.0248** | 1.1227<br>1.0000 | -0.6196<br>1.0000 | |
| **LL8** | -3.0150<br>**0.0128** | 0.3800<br>1.0000 | -0.7416<br>1.0000 | -0.0881<br>1.0000 |

**Dunn`s test with a Bonferroni correction: retinal layer (Z and p)**

| | GCL | INL |
|---|---|---|
| **INL** | 8.0511<br>**0.0000** | |
| **ONL** | 16.6914<br>**0.0000** | 8.6395<br>**0.0000** |

**b) Effect of light treatment on each retinal layer**

**<u>Ganglion Cell Layer (GCL)</u>**

**ANOVA test followed by Tukey's post hoc test**

| | **Df** | **Sum Sq** | **Mean Sq** | **F value** | **Pr (>F)** |
|---|---|---|---|---|---|
| **Light condition** | 4 | 65.73 | 16.433 | 9.136 | 2.11E-06 |
| **Residuals** | 109 | 196.05 | 1.799 | | |

| | **LD** | **LL2** | **LL4** | **LL6** |
|---|---|---|---|---|
| **LL2** | 1.60E-06 | | | |
| **LL4** | 0.03508 | 0.03436 | | |
| **LL6** | 0.00074 | 0.74849 | 0.57122 | |
| **LL8** | 0.00016 | 0.80203 | 0.39887 | 0.99984 |

Confidence level used: 0.95

**<u>Inner Nuclear Layer (INL)</u>**

**Kruskal-Wallis rank sum test**

| | **Chi-sq** | **dF** | **p** |
|---|---|---|---|
| **Light condition** | 22.84 | 4 | 0.000136 |

**Dunn`s test with a Bonferroni correction: light condition (p)**

| | **LD** | **LL2** | **LL4** | **LL6** |
|---|---|---|---|---|
| **LL2** | 0.0018 | | | |
| **LL4** | 0.0753 | 1.0000 | | |
| **LL6** | 0.0049 | 1.0000 | 1.0000 | |
| **LL8** | 0.0000 | 1.0000 | 0.1409 | 1.0000 |

Confidence level used: 0.95

**<u>Outer Nuclear Layer (ONL)</u>**

**Kruskal-Wallis rank sum test**

| | **Chi-sq** | **dF** | **p** |
|---|---|---|---|
| **Light condition** | 13.267 | 4 | 0.01004 |

**Dunn`s test with a Bonferroni correction: light condition (p)**

| | **LD** | **LL2** | **LL4** | **LL6** |
|---|---|---|---|---|
| **LL2** | 0.0128 | | | |
| **LL4** | 1.0000 | 0.0780 | | |
| **LL6** | 0.0600 | 1.0000 | 0.2727 | |
| **LL8** | 0.1680 | 1.0000 | 0.6952 | 1.0000 |

Confidence level used: 0.95

**c) Differential SG counts across retinal layers under different light treatments**

<u>**Light/Dark cycle (LD)**</u>

**Kruskal-Wallis rank sum test**

| | **Chi-sq** | **dF** | **p** |
|---|---|---|---|
| **Retinal layer** | 29.291 | 2 | 4.36E-07 |

**Dunn`s test with a Bonferroni correction: retinal layer (p)**

| | **GCL** | **INL** |
|---|---|---|
| **INL** | 0.0569 | |
| **ONL** | 0.0000 | 0.0016 |

<u>**Two days in continuous light (LL2)**</u>

**Kruskal-Wallis rank sum test**

| | **Chi-sq** | **dF** | **p** |
|---|---|---|---|
| **Retinal layer** | 63.497 | 2 | 1.63E-14 |

**Dunn`s test with a Bonferroni correction: retinal layer (p)**

| | **GCL** | **INL** |
|---|---|---|
| **INL** | 0.0001 | |
| **ONL** | 0.0000 | 0.0030 |

<u>**Four days in continuous light (LL4)**</u>

**Kruskal-Wallis rank sum test**

| | **Chi-sq** | **dF** | **p** |
|---|---|---|---|
| **Retinal layer** | 69.665 | 2 | 7.46E-16 |

**Dunn`s test with a Bonferroni correction: retinal layer (p)**

| | **GCL** | **INL** |
|---|---|---|
| **INL** | 0.0001 | |
| **ONL** | 0.0000 | 0.0000 |

Confidence level used: 0.95

<u>**Six days in continuous light (LL6)**</u>

**Kruskal-Wallis rank sum test**

| | **Chi-sq** | **dF** | **p** |
|---|---|---|---|
| **Retinal layer** | 53.645 | 2 | 2.24E-12 |

**Dunn`s test with a Bonferroni correction: retinal layer (p)**

| | **GCL** | **INL** |
|---|---|---|
| **INL** | 0.0001 | |
| **ONL** | 0.0000 | 0.0003 |

**Eight days in continuous light (LL8)**

**Kruskal-Wallis rank sum test**

| | Chi-sq | dF | p |
|---|---|---|---|
| **Retinal layer** | 66.692 | 2 | 3.30E-15 |

**Dunn`s test with a Bonferroni correction: retinal layer (p)**

| | GCL | INL |
|---|---|---|
| **INL** | 0.0002 | |
| **ONL** | 0.0000 | 0.0000 |
