## Supplemental material 2 for "Stress granule induction in rat retinas damaged by constant LED light"

### Supplemental Material 2: Normalization by area of the data presented in Figure 4

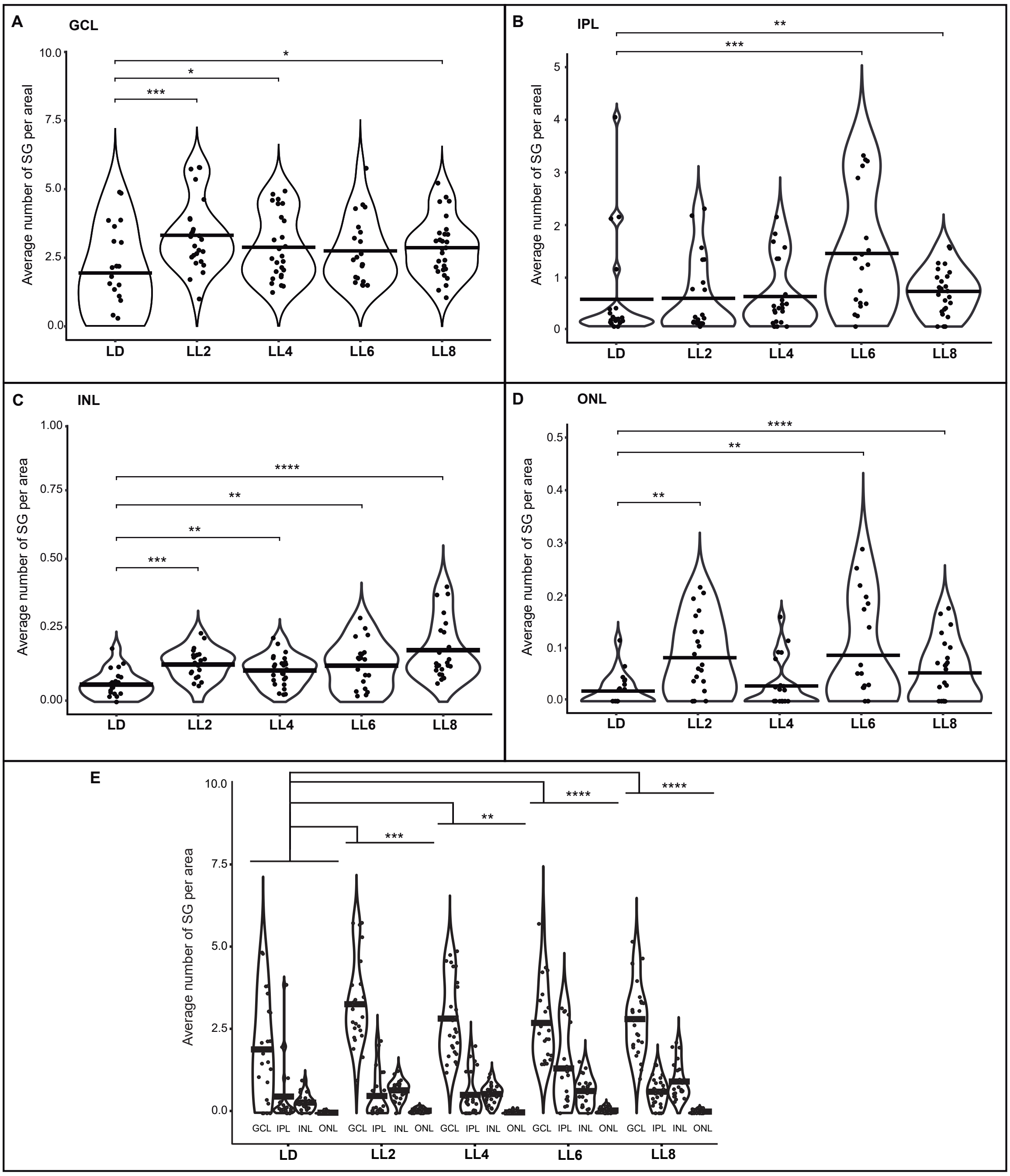

#### Statistical analysis

##### a) Combined effect of light exposure duration and retinal layer on SG count per area

**Scheirer-Ray-Hare test**

| | |
|---|---|
| **DV** | Number SG/Area |
| **Observations** | 462 |
| **D** | 0,99621 |
| **MS total** | 17825,5 |

| | Df | Sum Sq | H | p |
|---|---|---|---|---|
| **Light treatment** | 4 | 336958 | 18,975 | 0,00079 |
| **Retinal layer** | 3 | 5243822 | 295,294 | 0,00000 |
| **Light:Layer** | 12 | 186934 | 10,527 | 0,56985 |
| **Residuals** | 442 | 2407357 | | |

**Dunn`s test with a Bonferroni correction: light treatment (Z and p)**

| | LD | LL2 | LL4 | LL6 |
|---|---|---|---|---|
| **LL2** | -3,10088<br>**0,00096** | | | |
| **LL4** | -2,46831<br>**0,00679** | 0,69097<br>0,24479 | | |
| **LL6** | -3,79698<br>**0,00007** | -0,81426<br>0,20775 | -1,49199<br>0,06785 | |
| **LL8** | -3,79022<br>**0,00008** | -0,63213<br>0,26365 | -1,34965<br>0,08856 | 0,22253<br>0,41195 |

**Dunn`s test with a Bonferroni correction: retinal layer (Z and p)**

| | GCL | IPL | INL |
|---|---|---|---|
| **IPL** | 9,7652<br>**0,0000** | | |
| **INL** | 8,2734<br>**0,0000** | 1,4641<br>0,4295 | |
| **ONL** | 17,1058<br>**0,0000** | 7,1814<br>**0,0000** | 8,6487<br>**0,0000** |

##### b) Effect of light treatment on each retinal layer

**Ganglion Cell Layer (GCL)**

**Kruskal-Wallis rank sum test**

| | Chi-sq | dF | p |
|---|---|---|---|
| **Light condition** | 22,840 | 4 | 0,000054 |

| | LD | LL2 | LL4 | LL6 |
|---|---|---|---|---|
| **LL2** | 0,0004 | | | |
| **LL4** | 0,0178 | 0,0926 | | |
| **LL6** | 0,051 | 0,0522 | 0,3543 | |
| **LL8** | 0,0112 | 0,1158 | 0,4389 | 0,2999 |

Confidence level used: 0.95

**Inner Plexiform Layer (IPL)**

**Kruskal-Wallis rank sum test**

| | Chi-sq | dF | p |
|---|---|---|---|
| **Light condition** | 17,139 | 4 | 0,008170 |

**Dunn`s test with a Bonferroni correction: light condition (p)**

| | LD | LL2 | LL4 | LL6 |
|---|---|---|---|---|
| **LL2** | 0,2941 | | | |
| **LL4** | 0,1296 | 0,2826 | | |
| **LL6** | 0,0001 | 0,0008 | | |
| **LL8** | 0,0098 | 0,0384 | 0,1129 | 0,0595 |

Confidence level used: 0.95

**Iner Nuclear Layer (INL)**

**Kruskal-Wallis rank sum test**

| | Chi-sq | dF | p |
|---|---|---|---|
| **Light condition** | 25,117 | 4 | 0,000048 |

**Dunn`s test with a Bonferroni correction: light condition (p)**

| | LD | LL2 | LL4 | LL6 |
|---|---|---|---|---|
| **LL2** | 0,0001 | | | |
| **LL4** | 0,0065 | 0,0969 | | |
| **LL6** | 0,0038 | 0,1856 | 0,3681 | |
| **LL8** | 0,0000 | 0,1007 | 0,0039 | 0,0156 |

Confidence level used: 0.95

**Outer Nuclear Layer (ONL)**

**Kruskal-Wallis rank sum test**

| | Chi-sq | dF | p |
|---|---|---|---|
| **Light condition** | 15,121 | 4 | 0,00446 |

**Dunn`s test with a Bonferroni correction: light condition (p)**

| | LD | LL2 | LL4 | LL6 |
|---|---|---|---|---|
| **LL2** | 0,0011 | | | |
| **LL4** | 0,0649 | 0,0969 | | |
| **LL6** | 0,0038 | 0,1856 | 0,3681 | |
| **LL8** | 0,0000 | 0,1007 | 0,0039 | 0,0156 |

Confidence level used: 0.95

**c) Differential SG counts across retinal layers under different light treatments**

**Light/Dark cycle (LD)**

**Kruskal-Wallis rank sum test**

| | Chi-sq | dF | p |
|---|---|---|---|
| **Retinal layer** | 35,241 | 3 | 1,08E-07 |

**Dunn`s test with a Bonferroni correction: retinal layer (p)**

| | GCL | IPL | INL |
|---|---|---|---|
| **IPL** | 0,0027 | | |
| **INL** | 0,0223 | 0,2203 | |
| **ONL** | 0,0000 | 0,0010 | 0,0001 |

Confidence level used: 0.95

__Two days in continuous light (LL2)__

**Kruskal-Wallis rank sum test**

| | Chi-sq | dF | p |
|---|---|---|---|
| **Retinal layer** | 66,766 | 3 | 2,10E-14 |

**Dunn`s test with a Bonferroni correction: retinal layer (p)**

| | GCL | IPL | INL |
|---|---|---|---|
| **IPL** | 0,0000 | | |
| **INL** | 0,0000 | 0,0421 | |
| **ONL** | 0,0000 | 0,0246 | 0,0001 |

Confidence level used: 0.95

__Four days in continuous light (LL4)__

**Kruskal-Wallis rank sum test**

| | Chi-sq | dF | p |
|---|---|---|---|
| **Retinal layer** | 74,906 | 3 | 3,80E-16 |

**Dunn`s test with a Bonferroni correction: retinal layer (p)**

| | GCL | IPL | INL |
|---|---|---|---|
| **IPL** | 0,0000 | | |
| **INL** | 0,0000 | 0,1078 | |
| **ONL** | 0,0000 | 0,0010 | 0,0000 |

Confidence level used: 0.95

__Six days in continuous light (LL6)__

**Kruskal-Wallis rank sum test**

| | Chi-sq | dF | p |
|---|---|---|---|
| **Retinal layer** | 56,029 | 3 | 4,14E-12 |

**Dunn`s test with a Bonferroni correction: retinal layer (p)**

| | GCL | IPL | INL |
|---|---|---|---|
| **IPL** | 0,0027 | | |
| **INL** | 0,0000 | 0,0669 | |
| **ONL** | 0,0000 | 0,0000 | 0,0014 |

Confidence level used: 0.95

__Eight days in continuous light (LL8)__

**Kruskal-Wallis rank sum test**

| | Chi-sq | dF | p |
|---|---|---|---|
| **Retinal layer** | 79,935 | 3 | 2,20E-16 |

**Dunn`s test with a Bonferroni correction: retinal layer (p)**

| | GCL | IPL | INL |
|---|---|---|---|
| **IPL** | 0,0000 | | |
| **INL** | 0,0001 | 0,1135 | |
| **ONL** | 0,0000 | 0,0002 | 0,0000 |

Confidence level used: 0.95
